# Natural enemies of armored scales (Hemiptera: Diaspididae) and soft scales (Hemiptera: Coccoidae) in Chile: molecular and morphological identification

**DOI:** 10.1101/429357

**Authors:** P. Amouroux, D. Crochard, M.C.G. Correa, G. Groussier, P. Kreiter, C. Roman, E. Guerrieri, A.P. Garonna, T. Malausa, T. Zaviezo

## Abstract

Scale insects (Hemiptera: Coccoidea) are key pests of agricultural crops and ornamental plants worldwide. Their populations are difficult to control, even with insecticides, due to their cryptic habits. Moreover, there is growing concern over the use of synthetic pesticides for their control, due to deleterious environmental effects and the emergence of resistant populations of target pests. In this context, biological control may be an effective and sustainable approach. Hymenoptera Chalcidoidea includes natural enemies of scale insects that have been successfully used in many biological control programs. However, the correct identification of pest scale species and their natural enemies is particularly challenging because these insects are very small and highly specialized. Integrative taxonomy, coupling DNA barcoding and morphological analysis, has been successfully used to characterize pests and natural enemy species. In this study, we performed a survey of parasitoids and predators of armored and soft scales in Chile, based on 28S and COI barcodes. Fifty-three populations of Diaspididae and 79 populations of Coccidae were sampled over the entire length of the country, from Arica (18°S) to Frutillar (41°S), between January 2015 and February 2016. The phylogenetic relationships obtained by Bayesian inference from multilocus haplotypes revealed 40 putative species of Chalcidoidea, five Coccinellidae and three Neuroptera. In Chalcidoidea, 22 species were identified morphologically, resulting in new COI barcodes for 12 species and new 28S barcodes for 14 species. Two predator species (*Rhyzobius lophantae* and *Coccidophilus transandinus*) were identified morphologically, and two parasitoid species, *Chartocerus niger* and *Signiphora bifasciata*, were recorded for the first time in Chile.

## Introduction

Scale insects (Hemiptera: Coccoidea) are key pests of crops and ornamental plants worldwide. Diaspididae, Pseudococcidae, and Coccidae are the three most important families of Coccoidea, with 420, 259 and 169 genera, respectively (1). The control of these pests is still based essentially on repeated applications of synthetic insecticides, raising concerns about insecticide resistance in pests and possible effects on human health (2,3). A more sustainable approach to the management of these pests involves the use of resident natural enemies or the introduction of exotic ones (biological control), either alone or together with other control methods (integrated pest management) (4). The efficacy of biological control is dependent on the correct identification of both the target pest and its natural enemies (4). However, the morphological identification of scale insects requires considerable expertise, as it is based largely on microscopic cuticular characters visible only on adult females in most species. Genetic analyses have recently been added to the morphological approach for the integrative characterization of insects, and seem to be the only tool able to separate closely related species reliably (5,6). For example, DNA barcoding is a complementary tool for pest identification regardless of sex or developmental stage. This approach has recently been successfully applied to scale insects, with the molecular identification established for armored and soft scales in Chile (7) and for the mealybugs associated with grapes in Chile and elsewhere, worldwide (8–10).

The difficulties identifying scale insects also apply to the most important group of their natural enemies: Hymenoptera Chalcidoidea (11,12). The families Aphelinidae and Encyrtidae, in particular, have been successfully used in a number of biological control programs with Diaspididae and Coccoidea as target pests (13). However, fine-scale taxonomic identification is required for the implementation of appropriate control methods. The taxonomic diversity of pests and natural enemies may make it difficult to identify the most appropriate specific parasitoid (14), potentially leading to a failure to control the target pest (4). Correct identification also poses problems for predatory species from Chrysopidae (15), including *Chrysoperla* near *comanche* Banks, or Coccinellidae including *Rhyzobius lophantae* Blaisdell (16) and the *Coccidophilus* species commercially available for managing Chilean pest populations (17).

Since 1903, classical biological control programs involving the introduction of 87 biological control agents have been implemented in Chile (18). Eighteen of the species used targeted armored scales, including six species for *Aonidiella aurantii* Maskell, two for Lepidosaphes beckii (Newman), and one for *Comstockaspis perniciosa* (Comstock), or soft scales, including nine species for *Saissetia oleae* (Gómez-Menor Ortega). More than 50% of the biological control agents released became successfully established and now participate in the control of the targeted pest or closely related pest species (19). Biocontrol efficacy has also been enhanced by the activity of native parasitoid species, including *Aphytis chilensis* (Howard), *Aphytis proclia* (Walker), *Encarsia lycopersici*(De Santi), *Encarsia porteri* Mercet and *Coccidophilus citricola* Brethès, which attack armored scales, and *Coccophagus caridei* Brethès, which attacks soft scales (18,20). Chilean armored and soft scales have recently been characterized with molecular tools, but little effort has as yet been made to characterize their natural enemies and obtain barcodes for these species.

We present here the results of a survey of natural enemies (parasitoids and predators) attacking armored and soft scales in Chile, with the aim of providing basic information useful for future biological control programs. Our two objectives were: (i) to characterize the diversity of natural enemies of scale species sampled on various fruit crops, ornamental plants, and native trees and shrubs in Chile; and (ii) to describe the association between each scale species with its natural enemies in Chile.

## Materials and methods

### Sample collection

In total, 53 populations of Diaspididae were sampled from 36 sites and 79 populations of Coccidae were sampled from 48 sites along a latitudinal gradient from Arica in northern Chile (18°S) to Frutillar in southern Chile (41°S), between January 2015 and February 2016 (Fig. 1). Samples were collected from fruit crops, ornamental plants, and indigenous or endemic trees or shrubs. The minimum distance between two sampling sites was one kilometer. Each sample was checked and divided into subsamples. In each subsample, only one pest species was kept alive, the others being manually destroyed. Plant material for each subsample was placed in boxes and the emergence of natural enemies was monitored daily for 40 days. All emerging natural enemies were stored in 95% ethanol at –20°C until DNA extraction and/or morphological identification. All the emerging insects were sorted into morphospecies. Details of the samples collected (sampling locations, host plants, and dates) are available in S1 Table.

**Fig. 1:**
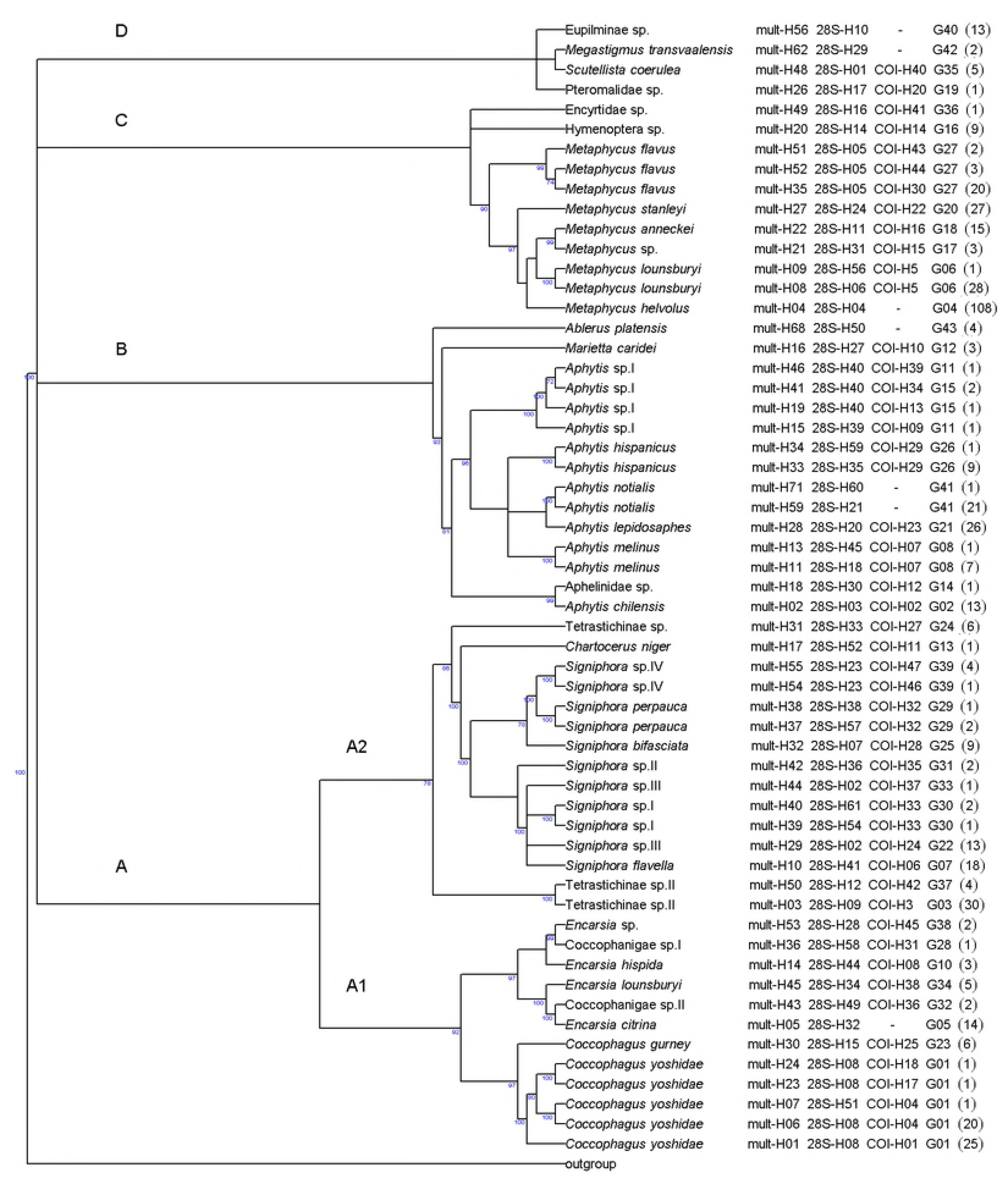
Sampling sites for Diaspididae (red upward-pointing triangle, *n*=36) and for Coccidae (blue downward-pointing triangle, *n*=48). Figure created with R software (21) (library ‘maps’(22)).

## DNA extraction and amplification

When available, two individuals per subsample and morphospecies were processed. DNA was extracted with the Prepgem Insect kit (Zygem, Hamilton, New Zealand). Parasitoids were not crushed before extraction and the time period over which the Prepgem enzyme was allowed to act was extended beyond the manufacturer’s recommendations (2 h at 75°C rather than 30 minutes). The total volume of 1x Prepgem Buffer and enzyme used was 30 μL per individual (23).

For each individual, we amplified two loci chosen on the basis of their suitability for DNA barcoding, population genetics and phylogenetic studies: the HCO-LCO region of the mitochondrial gene encoding cytochrome oxidase subunit I (COI) and the D2 region of the nuclear 28S gene (24). For COI we used the primers PCO-F1 (5’ CCTTCAACTAATCATAAAAATATYAG 3’) and Lep-R1 (5’ TAAACTTCTGGATGTCCAAAAAATCA 3’); and for 28S we used the primers C-28SLong-F (5’ GAGAGTTMAASAGTACGTGAAAC 3’) and C-28SLong-R (5’ TCGGARGGAACCAGCTACTA 3’). PCR was performed with a 23 μL reaction mixture and 2 μL of diluted DNA (1–20 ng). The reaction mixture contained 12.5 μL of 1x QIAGEN Multiplex PCR buffer and each of the primers required at a concentration of 0.2 μM. PCR was carried out as follows: initial denaturation at 95°C for 15 minutes, followed by 35 cycles of denaturation at 94°C for 30 s, annealing for 90 s at a temperature of 48°C (COI) or 58°C (28S), elongation at 72°C for 60 s, and then a final extension at 72°C for 10 minutes. The final products were separated by electrophoresis with the QIAxcel Advanced System (QIAGEN, Hilden, Germany) for quality control. The PCR products were then sent to Genewiz (UK, Essex), for bidirectional sequencing by capillary electrophoresis on an ABI 3130XL automatic sequencer (Applied Biosystems, Foster City, CA, USA). The consensus sequences were provided by Genewiz. Chromatograms were visualized with SeqTrace 0.9.0 (25) to check nucleotide variations. The sequences were deposited in GenBank with the following accession numbers: for the predators, MH455610 to MH455618 for COI and MH456371 to MH456401 for 28S; and for Chalcidoidea MH456402 to MH456789 for COI and MH455619 to MH456370 for 28S (S1 Table).

## Molecular identification and phylogenetic analysis

We considered sequences differing by one or more nucleotides to correspond to different haplotypes. Analyses were performed on the concatenated 28S and COI haplotypes (multilocus haplotype, abbr. mult-H). Regions of the 28S sequences displaying large numbers of insertions or deletions were removed for the analyses with Gblocks Server (26). Haplotype alignment was performed with MEGA version 7 (27) and the CLUSTALW method (28). Blast queries were performed against the NCBI GenBank database and final phylogenetic trees were generated from the haplotypes of each gene with R software (21): libraries “ape” (29), “ade4” (30), and “phangorn” (31). Species delineation was performed with the online version of Automatic Barcode Gap Discovery (ABGD) (32), with a prior maximal distance P=0.001 and a Kimura MinSlope distance of 1.0.

Bayesian inferences of phylogenetic relationships between multilocus haplotypes were obtained with BayesPhylogenies (33). Analyses were performed with nQ+C mixture models, with n varying between one and five independent rate matrices (Qs). The best model was chosen by comparing Bayes factors. We used a GTR model, as recommended by Pagel & Meade (2004). One Markov chain was used for ten million iterations and a print frequency of 1,000 iterations. The length of the burn-in period was determined by plotting likelihood across iterations. All iterations corresponding to the burn-in period (one million iterations) were removed from the output of BayesPhylogenies before subsequent analyses. We used the sump command of MrBayes (34) to obtain a summary of BayesPhylogenies outputs and to calculate Bayes factors. Majority-rule consensus trees were then drawn with PAUP*4 3.99 (35) (‘contree/Majrule’ command) from the output of the BayesPhylogenies analysis (9,000 trees), using the best model selected. Trees were rooted on concatenated *Mymaromma anomalum* (Blood and Kryger) (Hymenoptera: Mymarommatidae) sequences obtained from GenBank (accession numbers: GQ374772.1 for 28S and GQ374670.1 for COI). Mymarommatoidea was recently classified as a sister group of Chalcidoidea (36).

## Morphological identification

The morphological and molecular identification of scale insects is detailed a previous study by Amouroux et al. (7).

For the morphological characterization of parasitoids, adults from each haplotype with numerous individuals were selected, and mounted on cards and slides. For card mounting, parasitoids were transferred from absolute ethanol to a 1:1 ethanol: xylene solution for 24 h, then to amyl acetate for 24 h, and mounted on cards with water-soluble glue. For slide mounting, the wings were dissected from a card-mounted specimen and directly mounted in Canada balm on the slide (37). The remaining parts of the specimen were processed by incubation in 10% KOH for 5 min at 100°C, transferred to acetic acid for 5 minutes, then to a series of ethanol solutions of increasing concentration (from 70% to absolute) and finally to clove oil. The dissected parts of the insect (head, antennae, thorax, gaster, hypopygium, ovipositor, male genitalia) were mounted on the slide in Canada balm. The slide placed on a hot plate at 100°C for 2 h and Canada balm and cover slips were then placed over the dissected parts. For *Metaphycus*, the card- and slide-mounted specimens were compared with published descriptions (38,39) and authoritatively identified specimens deposited in the Natural History Museum of London (UK) collection. Aphelinidae and Signiphoridae were slide-mounted and compared with published descriptions (40–44) and authoritatively identified specimens deposited in the Museo di Entomologia Filippo Silvestri collection at Portici, Italy.

## Host-parasitoid associations

Sampling site was chosen as the unit for investigations of host-parasitoid associations. For each putative species, the percentage of Diaspididae (or Coccidae) at each sampling site was calculated to determine the dominant host family. We then determined the spatial distribution of each parasitoid, including all samples from the dominant host family. For specific host-parasitoid associations, we retained only associates observed at more than one sampling site.

## Results

### Molecular and morphological identification

#### Hymenoptera

DNA barcodes were successfully obtained for 763 hymenopteran individuals. For 28S, 731 sequences of 760 bp to 860 bp were obtained. Before Gblocks analysis to identify conserved blocks, the 28S sequences were reduced to 596 bp (without gaps) and assigned to 61 haplotypes. For COI, 386 sequences of 652 bp were obtained, corresponding to 50 haplotypes. COI sequences were shortened for only 10 individuals, with the removal of 5 bp (*n*=9) and 31 bp (*n*=1) from the start of the sequence. Both genes were successfully sequenced in 348 individuals. Following BLAST queries, we removed the sequences of individuals not belonging to Chalcidoidea. For subsequent analyses, we used data for 506 individuals, corresponding to 50 multilocus haplotypes and seven 28S haplotypes not associated with any COI haplotype.

Within the superfamily Chalcidoidea, ABGD analyses split the multilocus haplotypes into 42 groups, whereas the 28S sequences were split into 31 groups. By combining phylogenetic cladogram (Fig. 2) and ABGD group (Table 1) analysis, we identified 40 putative species. The intra variation of *Aphytis* sp. I and *Signiphora* sp. III was not maintained because both had specific 28S haplotype (Table 1) and the phylogenetic tree provided support for the monophyly of each cluster of haplotypes (Fig. 2). Morphological identification was possible for 22 of the 40 putative species. For the other 18 putative species, BLAST analyses of the GenBank database matched the DNA barcodes to a taxonomic reference. However, the identification retained was at a level above that of the species (genus or family), according to the phylogenetic cladogram. *Signiphora merceti* Malenotti was morphologically identified but was not associated with a barcoding sequence due to suspected cross-contamination. DNA barcodes were obtained for 28S and COI for 12 and 14 Chalcidoidea species without previous molecular characterization, respectively.

**Fig. 2:**
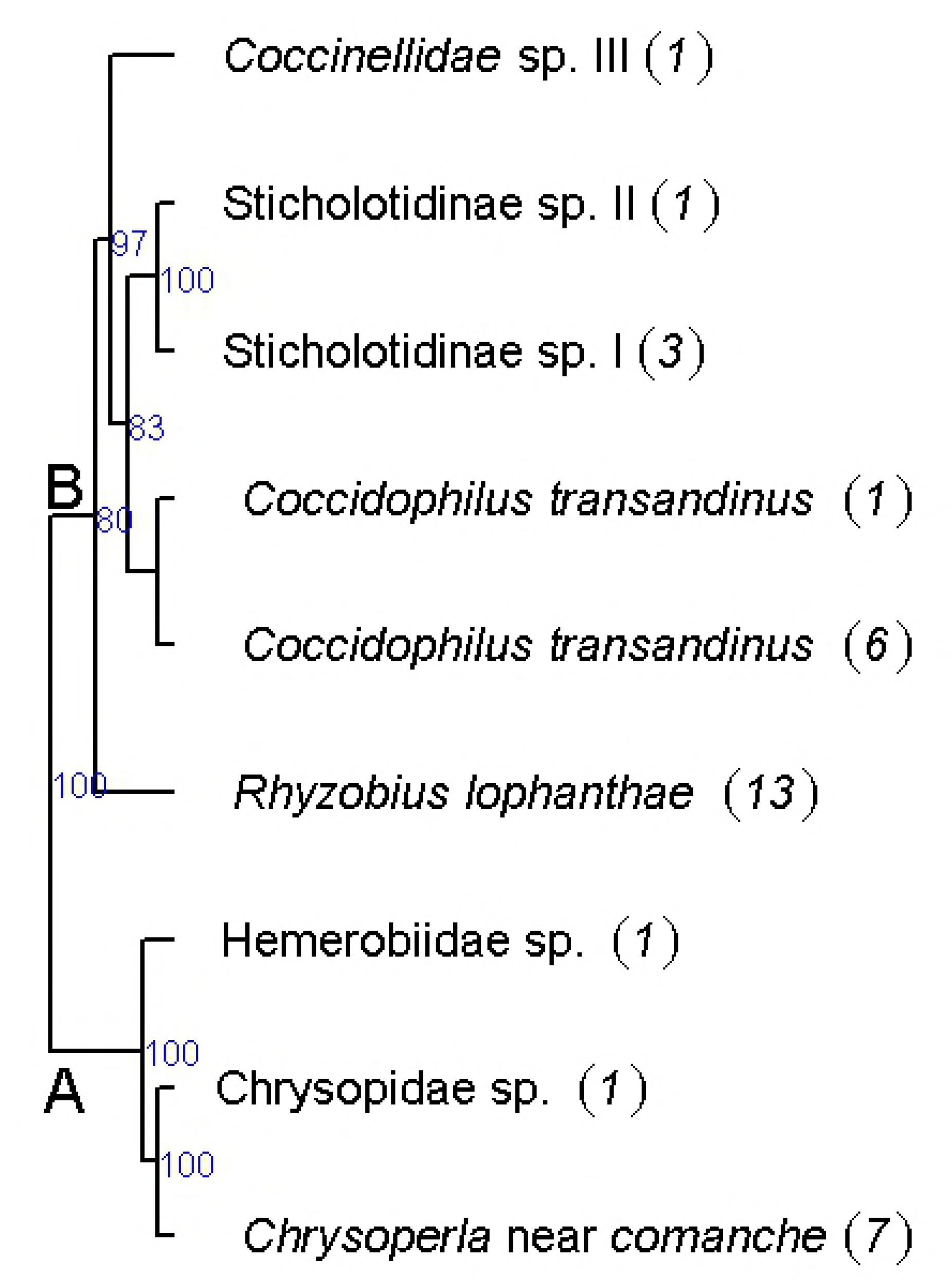
Phylogenetic tree of Chalcidoidea inferred with Bayesian methods (mixture models) from multilocus haplotypes. The majority-rule consensus tree was calculated from the Bayesian analysis. Bayesian posterior probabilities are represented beyond the nodes (values below 70% are not shown). Taxa are named according to their ABGD group, haplotype code and taxonomic names (Table 1).

**Table 1:**
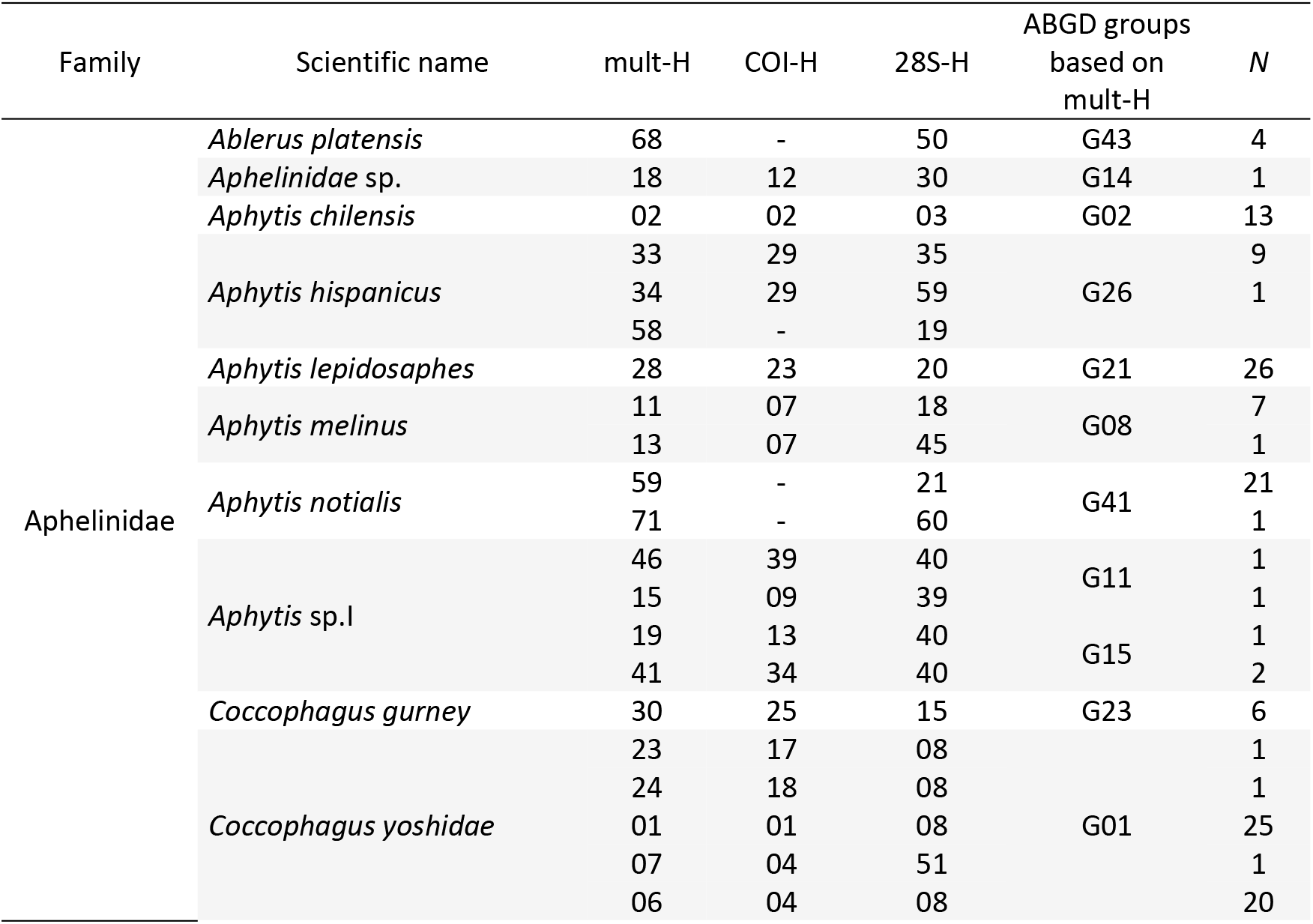
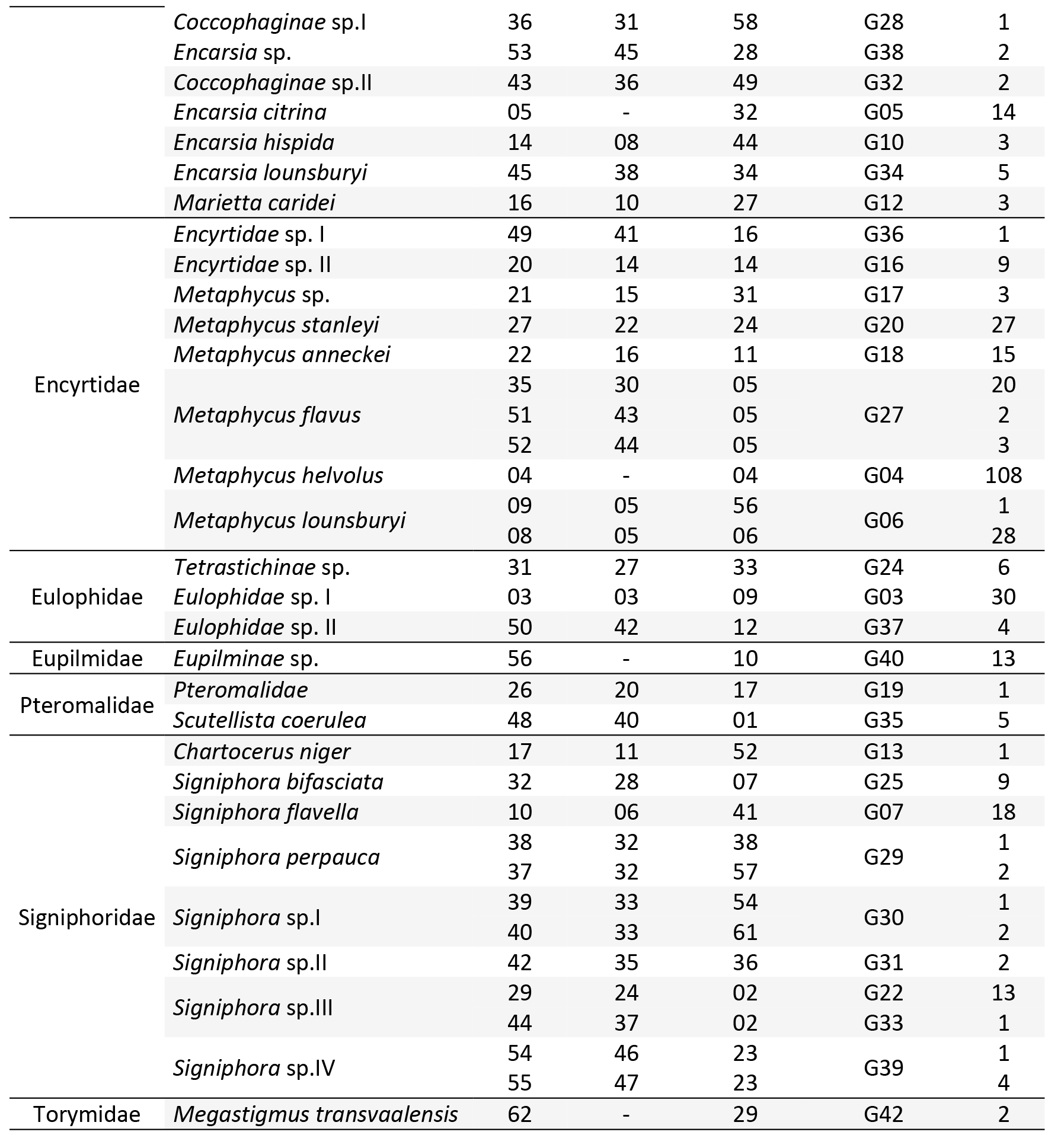
List of the 40 putative Chalcidoidea species found in the survey, with their haplotypes codes: multilocus haplotype (mult-H) resulting from concatenated 28S and COI sequences, COI haplotype (COI-H), and 28S haplotype (28S-H). ABGD groups correspond to putative species delineated by applying the ABGD method to multilocus haplotypes. *N* is the number of individuals for each multilocus haplotype.

**Table 2:**
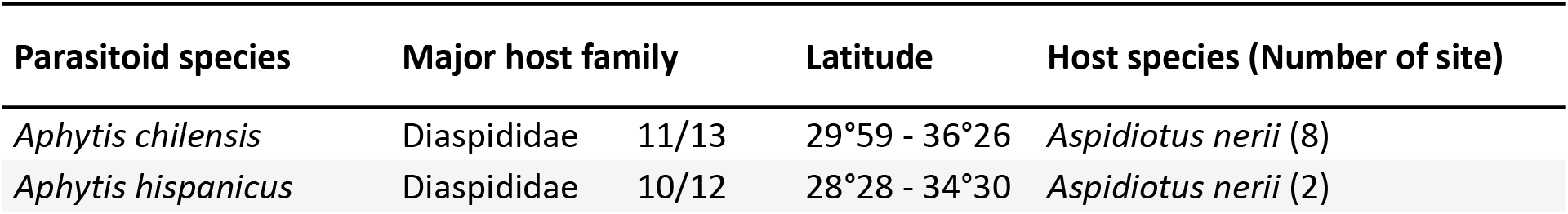
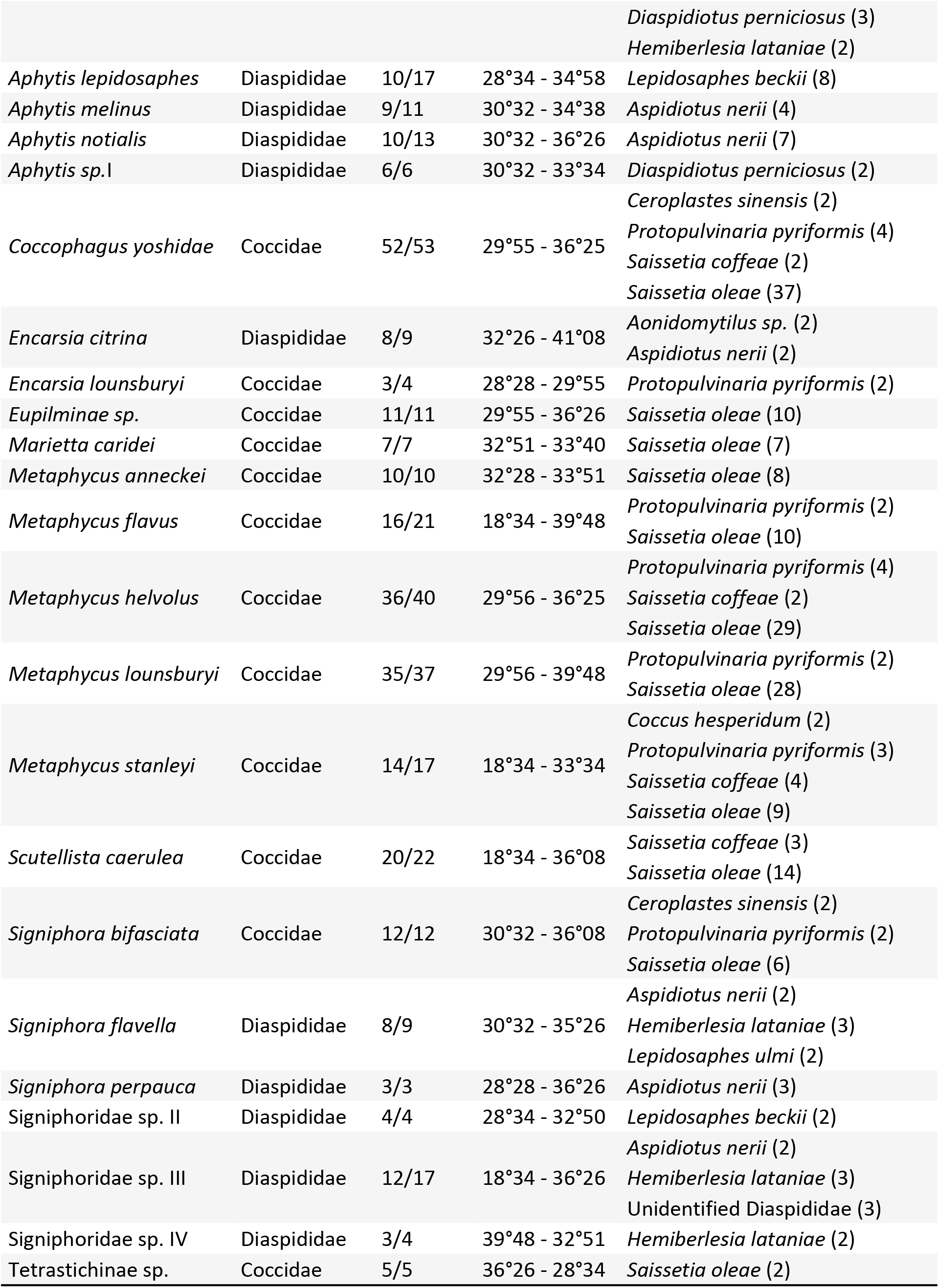
Chalcidoidea species, name of host family (number of samples from the majority host family / total number of samples), distribution (northernmost and southernmost latitudes), and list of host species (number of sites with the association)

On the cladogram based on multilocus haplotypes (Fig. 2), the family Encyrtidae (clade C) appeared to be monophyletic. The family Aphelinidae was found into two clades: A1 including the subfamily Coccophaginae, and clade B including the subfamily Aphelininae. Clade A2 contained the families Eulophidae and Signiphoridae. The families Pteromalidae, Torymidae and Eupelmidae were represented by only a few species and were grouped together in clade D.

#### Predators

DNA data were obtained for 34 individuals: 25 from Coleoptera Coccinellidae and nine from Neuroptera (Table 3). For 28S, 31 sequences of 764 bp to 986 bp were obtained. Before the Gblocks analysis, 28S sequences were reduced to 753 bp and five haplotypes were identified. For COI, eight sequences of 658 bp were obtained and seven haplotypes were identified. We considered nine haplotypes in the phylogenetic analysis: five haplotypes based on the two genes, three on 28S only and two on COI only.

**Table 3:** Predator species (Neuroptera or Coccinellidae) distribution (northernmost and southernmost latitudes), and number of individuals (*N)*.

| Predator species | Latitude | COI-P | 28S-P | *N* |
| --- | --- | --- | --- | --- |
| *Chrysoperla* near *comanche* | 32°49 - 34°38 | COI-P1 | 28S-P1 | 7 |
| Hemerobiidae sp. | 36°08 | - | 28S-P2 | 1 |
| Chrysopidae sp. | 32°51 | COI-P2 | - | 1 |
| *Rhyzobius lophanthae* | 28°28 - 34°38 | - | 28S-P3 | 13 |
| *Coccidophilus transandinus* | 32°26 - 33°51 | COI-P3 | 28S-P4 | 6 |
|  |  | COI-P4 | 28S-P4 | 1 |
| Sticholotidinae sp. I | 33°51 - 34°38 | COI-P5 | - | 3 |
| Sticholotidinae sp. II | 35°07 | COI-P6 | - | 1 |
| Coccinellidae sp. | 32°51 | COI-P7 | - | 1 |

Predators formed two clades (Fig. 3). Clade A contained three species from Neuroptera and clade B five species from Coccinellidae. *Chrysoperla* near *comanche* was identified morphologically by Bruno Michel (CIRAD – France) and *Coccidophilus transandinus* González was identified by Guillermo González (Chile).

**Fig. 3:**
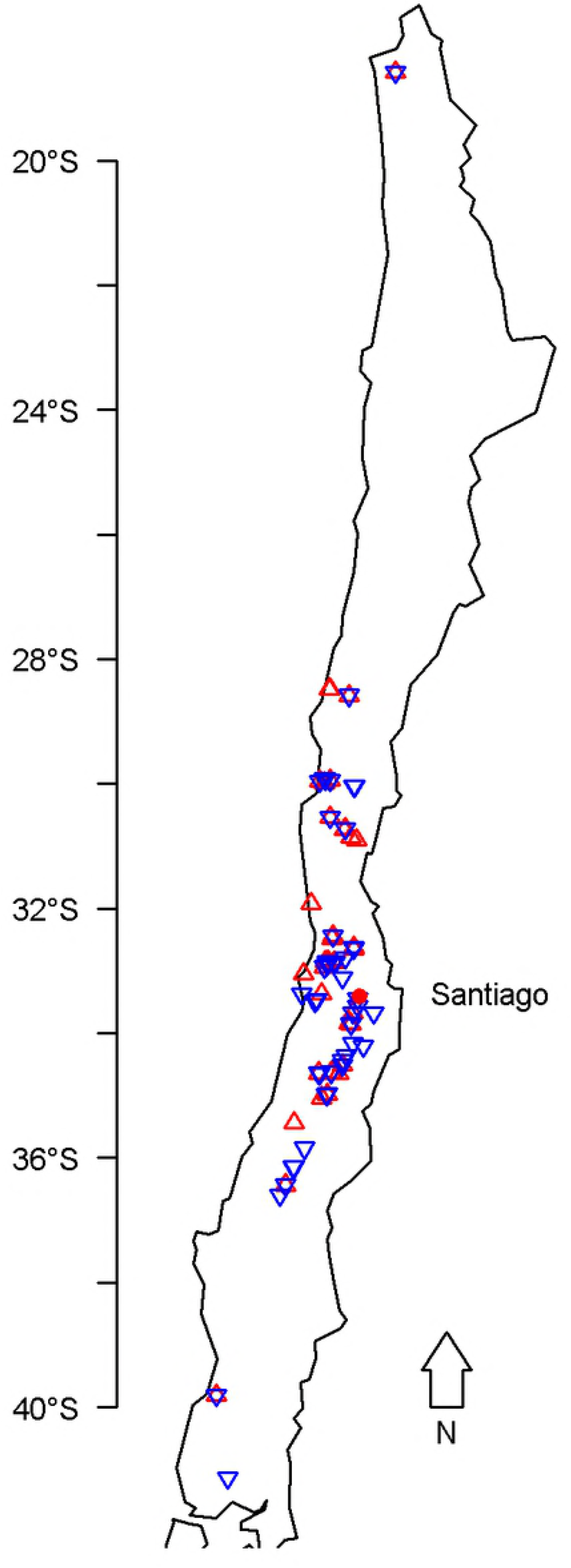
Phylogenetic tree of Neuroptera and Cocinellidae predators inferred by Bayesian methods (mixture models) from multilocus haplotypes. The majority-rule consensus tree was calculated from the Bayesian analysis. Bayesian posterior probabilities are shown beyond the nodes (values below 70% are not shown). Taxa are named according to their ABGD group, haplotype code and taxonomic names (Table 3).

### Host-parasitoid associations

Data for 738 DNA-barcoded individuals were used to develop a description of the geographic distribution of parasitoid species and hosts (Table 2).

Three species were discarded from this description: *Coccophagus gurneyi* Compere and *Encarsia hispida* De Santis, which are parasitoids of Pseudococcidae and Aleyrodidae, respectively; and *Megastigmus transvaalensis* (Hussey), which feeds on *Schinus* spp. seeds (13). Detailed information about all individuals is provided in S1 Table.

Armored scale insects from the surveyed sites were parasitized by 18 species from the families Aphelinidae, Azotidae and Signiphoridae. For the 13 species identified on the basis of morphology and DNA analyses, several previously unreported host associations were found, such as that between the apple mussel scale, *Lepidosaphes ulmi* (L.) and *Signiphora flavella* Girault.

Soft scale insects from the surveyed sites were parasitized by 12 putative species. For the nine Chalcidoidea species identified on the basis of morphology and DNA analyses, several previously unreported host associations were observed: *Coccophagus yoshidae* Peck on *Ceroplastes sinensis* Del Guercio, *Protopulvinaria pyriformis* Cockerell and *Saissetia coffeae* (Walker); *Signiphora bifasciata on C. sinensis, P. pyriformis* and *S. oleae*.

## Discussion

### Molecular and morphological identification, and phylogenetic analyses

In this survey, we obtained COI and 28S barcode sequences for 40 putative species from Chalcidoidea, five from Coccinellidae and three from Neuroptera from Chile. For 22 species of parasitoids, it was possible to assign a taxonomic name on the basis of morphological examination. One species, *Signiphora merceti*, was identified only by morphological examination. Two Signiphoridae species, *Chartocerus niger*(Ashmead) and *Signiphora bifasciata* Ashmead, have never before been reported in Chile. Thirteen (for COI) and 15 (for 28S) of the barcodes obtained are the first available for Chalcidoidea species and for predators.

Analyses based on multilocus haplotypes provided results that were consistent overall, with ABGD groups forming clusters on the phylogenetic trees. The only exception was *Aphytis* sp. I, for which the phylogenetic tree differentiated four groups, whereas ABGD analysis differentiated only two. However, the 28S sequences obtained were identical, so all individuals were retained in the same putative species.

One key finding was the division of the family Aphelinidae into two clades (A and B on Fig. 2), which also included Azotidae, Eulophidae and Signiphoridae. This division is congruent with the ‘eulophid lineage’ of Heraty et al. (36), which groups these these families of “soft-bodied” Chalcidoidea together. Moreover, these authors also reported that the family Aphelinidae was para- or polyphyletic. Before the work of Heraty et al. (34), Azotidae (including *Ablerus*) was considered to be a subfamily of Aphelinidae (= Azotinae)(45).

For predators, the two main clades identified were consistent with insect orders, with a single branch containing three species of Neuroptera, including one species of Hemerobiidae and two species of Chrysopidae, and another branch containing five species of Coccinellidae (Coleoptera), including one species of Coccidulinae (*R. lophanthae*) and three species of Sticholotidinae including *C. transandinus*.

### Species of natural enemies attacking scale insects in Chile

Aphelinidae displayed the greatest diversity, with seven species in the subfamily Aphelininae, including six identified to species level: *Aphytis lepidosaphes* (Mercet), *Aphytis melinus* De Bach, *Aphytis notialis De Santis, Aphytis hispanicus* (Mercet), *A. chilensis* and, *Marietta caridei Brèthes. Seven species from Coccophaginae were collected, five of which were identified* to species level: *Coccophagus gurneyi, C. yoshidae, Encarsia citrina* (Crawford), *E. lounsburyi*(Berlese & Paoli), *E. hispida*. In total, 11 species of *Aphytis* are known in Chile (13), most of which were introduced for classical biological control (18). *A. melinus* (introduced in 1966) and *A. lepidosaphes* (introduced in 1951) were the only introduced species recovered in this survey. We also collected the native species *A. chilensis*, which is used in biological control and has been introduced into other countries from Chile (19). The other *Aphytis* taxon collected in this study, *Aphytis* sp. I, may represent a new species.

Within the subfamily Coccophaginae, *Encarsia lounsburyi* was observed on various Coccidae species. This species was widely distributed throughout Chile and had a high degree of genetic diversity. Only *E. citrina* was collected on Diaspididae. *Coccophagus lycimnia* (Förster) and *C. caridei*, two members of Coccophaginae widely considered to be natural enemies of Coccidae species in Chile (46), were not recovered in our survey.

The family Signiphoridae was represented by at least eight species, including *Signiphora perpauca* Girault, *S. flavella, S. merceti, S. bifasciata* and *Chartocerus niger*. The last two of these species have never before been recorded in Chile. *Signiphora aspidioti* Ashmead and unidentified Signiphoridae species have been reported as natural enemies of *Hemiberlesia* spp. in Chile (46). However, none of these species has yet been used in augmentative biological control programs in Chile (18,19). Given their small size, accidental introduction and the existence of native species cannot be ruled out. Wooley and Dal Molin (44) highlighted the importance of *Signiphora* as primary parasitoids of armored scales, including *S. perpauca* against *Chrysomphalus aonidum* (L.) in Brazil. In this study, *S. perpauca* was associated only with *Aspidiotus nerii* Bouché. The other six species of *Signiphora* were collected on Diaspididae (*Hemiberlesia lataniae*(Signoret), *A. nerii*, and *Lepidosaphes* spp.), whereas only *S. bifasciata* was found on Coccoidea (Table 2). *S. bifasciata* has been reported to be a hyperparasitoid of Coccidae in Mexico (43). Signiphoridae may be important natural enemies of scale insects in Chile and probably deserve more attention.

In our survey, Encyrtidae was represented by eight putative species, including six *Metaphycus* species: *Metaphycus anneckei* Guerrieri & Noyes*, Metaphycus flavus* (Howard)*, Metaphycus helvolus, Metaphycus stanleyi*(Compere), *M. lounsburyi*, and one unidentified species. *Metaphycus flavus* is a cosmopolitan species present in Chile. The other four species were all introduced for classical biological control of *S. oleae* between 1931 and 1997. They now all contribute to the natural control of other Coccidae species, including *Coccus hesperidium* L., *S. coffeae*, and *P. pyriformis*. Ripa & Larral (46) identified the same five species of parasitoids and an undescribed species attacking first to third instars of *S. oleae*. However, the number of individuals collected (*n*=3) was too small for further characterization of this taxon.

*Scutellista caerulea (Fonscolombe) is one of the two species from Pteromalidae collected. It was* introduced into Chile in 1931 and is now present over a wide range of latitudes. It is an efficient biocontrol agent against Coccidae due to its behavior as a predator of eggs or as an ectoparasitoid of adults (46). Its activity complements that of other *Metaphycus* spp., which attack young larval stages of the same host.

The predators identified in this study included three putative species of Neuroptera, two from Chrysopidae and from Hemerobiidae. Only *Chrysoperla* near *comanche* was identified morphologically. Molecular identification within Chrysopidae was inconclusive, probably due to the occurrence of cryptic species, as observed in the Northern Hemisphere and Africa (47). The larvae of these species are voracious predators known to be generalists but including scale insects among their prey (46).

Five Coccinellidae species were collected. *Coccidophilus transandinus* (Sticholotidinae) was identified morphologically, whereas *Rhyzobius lophantae* (Coccinellidae: Coccidulinae) was clearly identified on the basis of similarities between its DNA sequences and sequences present in the GenBank database. This common biocontrol agent was introduced into Chile in 1931 for the control of *Aonidiella aurantii* (18), and is now available commercially. The other three species belonged to the subfamily Sticholotidinae, which includes 17 species reported to be present in Chile, all belonging to tribe Microweiseini (48). Their small size makes them difficult to collect, but they are considered to be efficient natural enemies of armored scale insects and are reared for this purpose (48).

## Conclusion

We report COI and 28S barcode sequences for parasitoids and predators of armored and soft scales from Chile, which will facilitate the identification of these species in the future. Two Signiphoridae species new to Chile were recorded, and potential new species were detected. Host-parasitoid associations merit greater attention, to improve our understanding of the potential role of some species in the biological control of insect pests. Further research could focus on little studied groups, such as Signiphoridae, and on natural enemies of native scale insects. A deeper knowledge of the biology of the most promising biological control agents is required, together with additional information about possible interactions between species attacking the same host.

## Acknowlegments

We would like to thank all the growers who allowed us access to their orchards for sample collection. We thank Paula Molina and Angela Romo for technical assistance in the laboratory. We also thank all those who helped us to collect samples throughout Chile: Hugo Herrera, Francisco Bugueño, Sebastian Catalan and Isaac Vera from Xilema-ANASAC (Quillota); Claudio Salas from Instituto de Investigaciones Agropecuarias (INIA Intihuasi – La Serena); Marta Godoy from Cabilfrut (Ovalle); Leonardo Fuentes from Pontificia Universidad Católica de Valparaíso (Quillota); Alda Romero from Pontificia Universidad Católica de Chile (Santiago); Jean-Claude Malausa from Institut National de la Recherche Agronomique (Sophia-Antipolis, France). This project was supported by the EU FP7 Marie Curie IAPP Project “Colbics” #324475, the “Fundación para la Innovación Agraria” (FIA) #PYT-2015–0230 and CONICYT FONDECYT Postdoctorado No. 3180344

## Supporting information

**S1 Table.** List, in alphabetical order, of the 798 individuals sequenced, their molecular code for the voucher in the INRA collection, their sample date and location (administrative region, municipality, longitude and latitude, elevation), their host plant (scientific name), their family and species host (scientific name), and their number Genbank accession number for COI and 28S.

